# Pharmacological inhibition of LIN28A promotes imatinib sensitivity in CML resistance

**DOI:** 10.64898/2026.04.27.720799

**Authors:** Owen F.J. Hovey, Alyssa Wu, Tingting Wu, Jenica H. Kakadia, Mallory I. Frederick, Ilka U. Heinemann, Shawn S. C. Li

**Author notes:** Corresponding author: Shawn S.C. Li **Email:**.

## Abstract

Resistance to tyrosine kinase inhibitors (TKIs) remains a critical challenge in chronic myeloid leukemia (CML), particularly when driven by mechanisms independent of BCR-ABL1 kinase-domain mutations. Building on the identification of the RNA-binding protein LIN28A as a driver of imatinib resistance, we evaluated emerging LIN28 inhibitors as potential sensitizing agents. Screening three small molecules in an imatinib-resistant (ImR) K562 model identified LIN28i-1632 as uniquely synergistic with imatinib (synergy score: 12.07), reducing cell proliferation by 71.15%. Quantitative DIA and TMT proteomics revealed that this synergy is characterized by significant proteomic remodelling, including the downregulation of the canonical LIN28 target HMGA1 and the activation of apoptotic and G2/M cell cycle checkpoint programs. Mechanistically, phosphoproteome and kinome profiling showed suppressed AKT/RPS6K and CDK signalling. We further demonstrate that LIN28i-1632 promotes sensitivity by reducing BCR-ABL protein abundance and attenuating the AKT survival axis through RICTOR downregulation and PTEN restoration. Collectively, our findings establish pharmacological LIN28 inhibition as a viable strategy to overcome TKI resistance by simultaneously engaging cell-cycle arrest and dismantling the AKT-mediated survival network.

## Introduction

Each year, chronic myeloid leukemia (CML) accounts for 15% of new leukemia cases in the United States ^1^. CML is caused by the translocation of chromosomes 9 and 22, forming what is known as the Philadelphia (Ph) chromosome and producing the fusion protein BCR-ABL from the breakpoint cluster region (BCR) and the tyrosine-protein kinase ABL1 ^2^. While multiple generations of tyrosine kinase inhibitors (TKIs), including imatinib, dasatinib, nilotinib, bosutinib, and ponatinib ^3^, have been developed to target the BCR-ABL protein, imatinib remains a primary treatment despite rising resistance ^4,5^. While many patients remain on TKIs for life, approximately 20-30% achieve treatment-free remission ^6^. CML progresses through the chronic phase, the accelerated phase, and finally to the blast crisis, with each phase having a higher proportion of blast cells. CML becomes more resistant to TKIs as patients progress through the stages ^7^.

Two RNA-binding proteins, LIN28A and LIN28B, are important regulators of pluripotency and glucose metabolism ^8^. To regulate cell fate, LIN28A/B employs both let-7-dependent degradation—via TUT4 recruitment and uridylation—and let-7-independent translation enhancement, achieved by directly binding mRNAs like Igf2 to stabilize their association with the translation initiation complex ^9^. High expression of LIN28A/B are found in many cancers, including breast, lung, ovarian, brain, and acute myeloid leukemia, which is associated with poor prognosis ^9^.

Previously, we showed that LIN28A promoted resistance in CML through rewiring of translation and the kinome^10^. Overexpression of LIN28A activated multiple pathways, including AKT, PKC, SGK, and RPS6K, thereby promoting survival and proliferation. LIN28A/B has previously been detected in 40% of patients with advanced stages of CML ^11^. Furthermore, when paired CML patient samples from the chronic phase and blast crisis were compared, 23% of patients showed increased LIN28B after progression to blast crisis.

Recent work has now identified multiple LIN28 inhibitors including LIN28i-1632, LI71, SB1301, C902, PH-43, TPEN, Ln7, Ln15, and Ln115 ^12-14^. In 2016, 16,000 small molecules were screened using a FRET based sensor with LIN28B and let-7 which identified LIN28i-1632 as the most promising inhibitor of LIN28 activity ^14^. In 2018, 101,017 compounds were screened using fluorescent polarization with a pre-let-7f probe, identifying TPEN and LI71 as LIN28A/B inhibitors ^13^. Most recently, a study screened 18 million *in silico*, of which 163 were screened by fluorescent polarization, selecting the top three compounds Ln7, Ln15, and Ln115 ^12^. LIN28i-1632 and LI-71 were screened and shown to block ovarian and prostate cancer stemness phenotypes ^12^. LIN28i-1632 has also been shown to inhibit ovarian cancer growth through FAK/PTK2 by the LIN28B-let-7 axis ^15^. Treatment with LI-71 inhibited LIN28 function, increasing let-7g expression without affecting LIN28A-let-7-independent functions in the YAP1 pathway ^16^. This indicates that LI-71 may only be effective on the LIN28-let-7 pathway and not in LIN28A alternative functions. Additionally, in murine models, LIN28i-1632 has been shown to protect against non-alcoholic fatty liver disease by targeting NCoR1 through both the mTOR pathway and independently of the mTOR pathway ^17^. Targeting LIN28, based on these initial studies using LIN28 inhibitors, shows promise as a potential approach to combating cancer stem cells and resistant cancer.

Building on prior work implicating LIN28 in TKI resistance in CML^10,18,19^, we demonstrate that among three recently developed inhibitors, LIN28i-1632 uniquely synergizes with imatinib. This combination overcomes LIN28A-mediated resistance by inducing apoptosis, modulating cell cycle signalling, and suppressing the AKT pathway.

## Methods

### Cell Culture

ImR cells derived from K562 by our lab^10^ and were maintained in Iscove’s Modified Dulbecco’s Medium (IMDM) supplemented with GlutaMAX™ (Thermo Fisher Scientific, 31980030) and 10% fetal bovine serum (Wisent, 098-150) at 37°C in a humidified 5% CO_2_ atmosphere. Cells were treated with inhibitors, including imatinib (Selleckchem; S2475), TPEN (MedChemExpress; HY-100202), LIN28i-1632 (Toris; 6068), LI71 (MedChemExpress; HY-123905).

### Cell Viability Assays

Relative cell number was measured by WST-8 Cell Proliferation Assay Kit (Cayman Chemical; 10010199). Cells were plated at 100,000 cells/mL with increasing concentrations of inhibitors. After 72 hours, 10 μL of WST-8 reagent was added and incubated for 30 minutes at 37°C, 5% CO_2_. Optical density (OD) was measured at 450 nm, and background subtraction was performed at 650 nm. The data was analyzed with SynergyFinder 2.0 to identify ideal concentrations ^20^.

### RT-qPCR

Reverse transcription and quantitative PCR (RT-qPCR) of miRNAs and mRNAs were performed as described ^21^. Briefly, the total RNA, including miRNAs, was extracted from cell pellets using Qiagen miRNeasy Tissue/Cells Advanced Mini Kit (Qiagen, 217604) and quantified on Thermo Fisher Nanodrop 2000 spectrophotometer. Total RNA (2 μg) was reverse transcribed using the High-Capacity cDNA Reverse Transcription Kit (Applied Biosystems, 4368814) with miRNA-specific primers for miRNA targets and random primers for mRNAs. qPCR was performed on CFX connect (Bio-Rad) or Quantstudio 3 (Applied Biosystems) using PowerTrack SYBR Green (Applied Biosystems; A46110) and target-specific primers. Results were quantified by the delta-delta-CT method ^22^.

### Immunoblotting

Cells were lysed in NP-40 buffer (1% NP-40, 1% SDS, 20 mM Tris pH 7.6, 150 mM NaCl, 100 mM NaF, 1 mM PSMF, 1x Phosphatase inhibitor cocktail) and pulse sonicated for 5 seconds for a total of 20 seconds at 35% amplitude, then centrifuged at 20,000×g for 10 minutes. Protein concentration was measured using the Pierce™ BCA Protein Assay Kit (Thermo Scientific; 23227). 20 μg of sample with 1x Laemmli buffer (BioRad; 161-0747) was heated at 95°C for 5 minutes. Extracted proteins were run on either 8% or 12% SDS-page gel depending on the protein size and transferred to Immobilon® -PSQ PVDF Membrane (Millipore Sigma; ISEQ00010) at 30 V overnight at 4°C. Membranes were stained with FCF green solution and washed with 6.7% acetic acid and 30% methanol. Total protein was imaged on a ChemiDoc™ MP imager (BioRad) using the IR680 channel. Blots were blocked with 2% fish gelatin in 0.1% TBST for 45 minutes at room temperature. Primary antibodies were incubated overnight with shaking at 4°C. Blots were washed five times for 5 minutes with 0.1% TBST. Secondary antibodies were incubated for 45 minutes with either StarBright Blue 700 Goat Anti-Mouse IgG (Bio-Rad; 12004158), Donkey anti-Goat IgG (H+L) Highly Cross-Adsorbed Secondary Antibody, Alexa Fluor™ Plus 680 (Invitrogen; A32860) or Donkey anti-Rabbit IgG (H+L) Highly Cross-Adsorbed Secondary Antibody, Alexa Fluor™ Plus 800 (Invitrogen; A32808). Blots were imaged using ChemiDoc™ MP with either the Starbright 700, IR680 or IR800 channels. Quantification was performed in ImageLab (Bio-Rad; version 6.1.0) with the rolling circle method and normalized to total protein.

### Mass Spectrometry Sample Preparation

For data-independent acquisition (DIA), 20 μg of protein was reduced and alkylated with 5 mM TCEP (BioShop; 51805-45-9) and 15 mM IAA (Sigma-Aldrich; I1149), respectively^23^. The samples were cleaned up with Sera-Mag™ Carboxylate-Modified Magnetic Beads & SpeedBeads [E3 and E7] (Cytiva; 44152105050250 and 24152105050250) using the SP3 protocol and resuspended in 50 mM ammonium bicarbonate (ABC) pH 8, as previously described. Samples were digested with 1.5 mAU of LysC (Wako; 129-02541) for 2 hours, followed by Trypsin (Promega; V5113) at a 1:50 ratio at 37 °C for 16 hours. Peptides were desalted with C18 stage tips and quantified (A280) using a Take3 plate on a Cytation 1 imaging reader to a final concentration of 100 ng/μL.

For TMT-labelled samples, 150 µg of protein was reduced and alkylated as described above. Samples were cleaned using SP3 beads, resuspended in 50 mM EPPS pH 8.5 (Sigma-Aldrich). Following LysC and Trypsin digestion, peptide concentrations were determined using a BCA assay. From each sample, 100 μg of peptides were labelled with TMT11 isobaric tags (Thermo Fisher Scientific; 90110) and pooled. A 5% aliquot of the pooled sample was fractionated using the Pierce™ High pH Reversed-Phase Kit (Thermo Fisher Scientific; 84868). Phosphopeptides were enriched using the High Select™ TiO_2_ Phosphopeptide Enrichment Kit (Thermo Fisher Scientific; 88303) prior to High pH fractionation according to the manufacturer’s instructions.

### LC-MS/MS Acquisition

DIA samples were analyzed on an UltiMate™ 3000 RSLCnano system (Thermo Fisher Scientific) coupled to a Q-Exactive Plus Mass Spectrometer using an EASY-Spray source (Thermo Fisher Scientific). Mobile phases consisted of 0.5% acetic acid in water (Buffer A) and 0.5% acetic acid in 80% acetonitrile (Buffer B). Peptides were trapped on a PepMap Neo Trap Cartridge (5 mm × 300 μm, Thermo Fisher Scientific; 174500) and separated on an EasySpray column (2 μm C18, 75 μm × 250 mm, Thermo Fisher Scientific; ES902) at a flow rate of 300 nL/min. The DIA method comprised of a full MS scan followed by 19 windows (20 m/z width, 1 m/z overlap) covering 400-837 at a resolution of 17,500 (AGC target 5e5). A chromatogram library was generated using gas-phase fractionation using seven staggered m/z ranges with 4 m/z windows.

TMT samples were analyzed by data-dependent acquisition (DDA) as previously described^23^. Peptides were separated on a Double nanoViper PepMap Neo column (2 μm C18, 75 μm × 500 mm, Thermo Fisher Scientific; DNV75500PN). Separation was performed at 300 nL/min for total proteome (4-hour gradient) and pST (3-hour gradient) injections.

#### MS Data Processing

Raw DIA files were demultiplexed and peak-picked using MSConvertGUI ^24^. A gas-phase fractionation library was generated in Fragpipe (v20.0) using the *Homo sapiens* reference proteome (UP000005640; 20,460 entries). DIA-NN (v1.8.1) was used for searching with a 1% FDR filter and a precursor range of 400-837 m/z.

TMT data were processed using Fragpipe (v20.0)^25^ with default TMT11-bridge (proteome) and TMT11-phospho-bridge (pS/pT) workflows. Search parameters allowed for two missed cleavages and a mass tolerance of 20 ppm for precursors and fragments. Fixed modifications included carbamidomethylation (+57.02146) on cysteine and TMT (+229.16293) on lysine. Variable modifications included methionine oxidation (+15.9949), N-terminal acetylation (+42.0106), and phosphorylation (+79.96633) for the phospho-workflow. Phosphosite localization was filtered for Class I sites (probability ≥ 0.75). Peptide-spectrum matches (PSMs) were filtered to 1% FDR using Percolator, and TMT-integrator was used for quantification, excluding all peptides with missed cleavages.

### Bioinformatics Analysis

Proteomic data were processed and analyzed using R (v4.2.2). Initial data refinement included filtering for proteins present in all replicates of at least one sample type, followed by missing value imputation using the RandomForest algorithm. DIA samples were normalized using the Loess regression. For differential expression analysis, *DEqMS* ^26^ was used for total proteome calculations, while *limma*^27^ was used for phosphoproteomic data; both utilized the

Benjamini-Hochberg correction to control the False Discovery Rate (FDR). Functional enrichment was performed using gene set enrichment analysis (GSEA)^28^ for the proteome and kinase substrate enrichment analysis (KSEA)^29^ with PhosphoSitePlus^30^ data downloaded on August 22, 2023 for the phosphoproteome. Finally, protein interaction networks were mapped in Cytoscape (v3.8.0)^31^ and visualized using OmicsVisualizer (version 1.3.1)^32^ .

Data analysis and visualization were performed using GraphPad Prism (v9.5.1) or R (v.4.1.2) with the *ggplot2* package. Unless otherwise specified, data are presented as mean ± SEM from at least three independent biological replicates (n ≥ 3). Statistical significance was determined using Student’s t-tests for two-group comparisons or one-way ANOVA followed by Tukey’s post-hoc test for multiple comparisons. P-values are represented as follows: *P < 0.05, **P < 0.01, ***P < 0.001, ****P < 0.0001; *ns* indicates non-significance (P ≥ 0.05).

## Results

### LIN28i-1632 Uniquely Sensitizes ImR Cells to Imatinib

Building on our identification of LIN28A as a driver of imatinib resistance, we screened three commercially available LIN28 inhibitors – LIK-71, TPEN, and LIN28i-1632 – for synergistic effects with imatinib in ImR cells (Figure 1).

**Figure 1:**
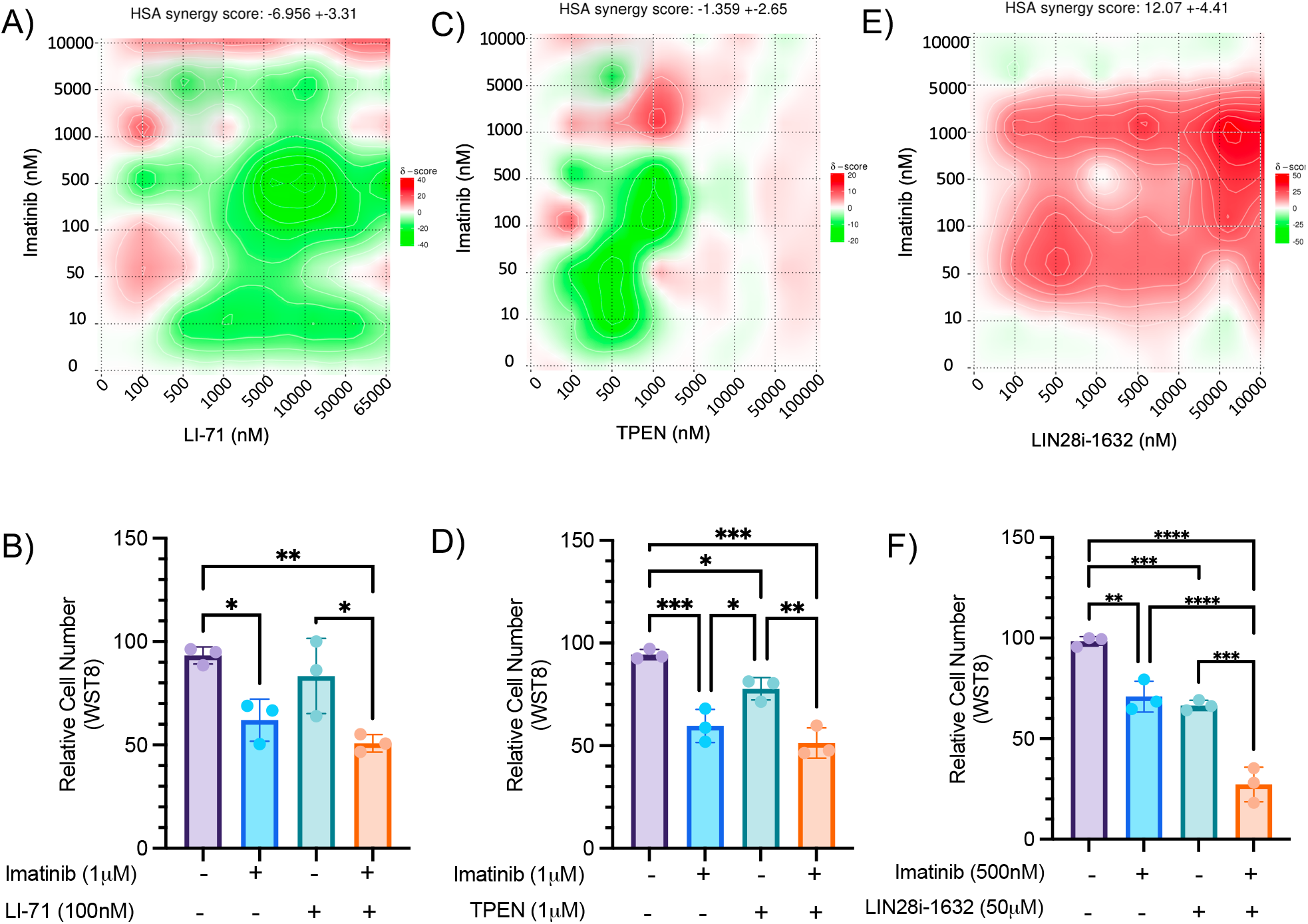
Screening of LIN28A inhibitors in combination with imatinib in resistance. The relative cell number across concentrations of imatinib from 10nM to 10 μM and either LI-71 (A), TPEN (C), or LIN28i-1632 (E), using SynergyFinder to identify synergistic regions. The most synergistic regions with imatinib presented for each of the combinations with LI-71 at 100nM (B), TPEN at 1μM (C) and LIN28i-1632 at (F). Values were graphed as mean±SEM and statistical significance was determined by ANOVA test: *P <0.05; **P < 0.01; ***P < 0.001; ****P < 0.0001.

**Figure 2:**
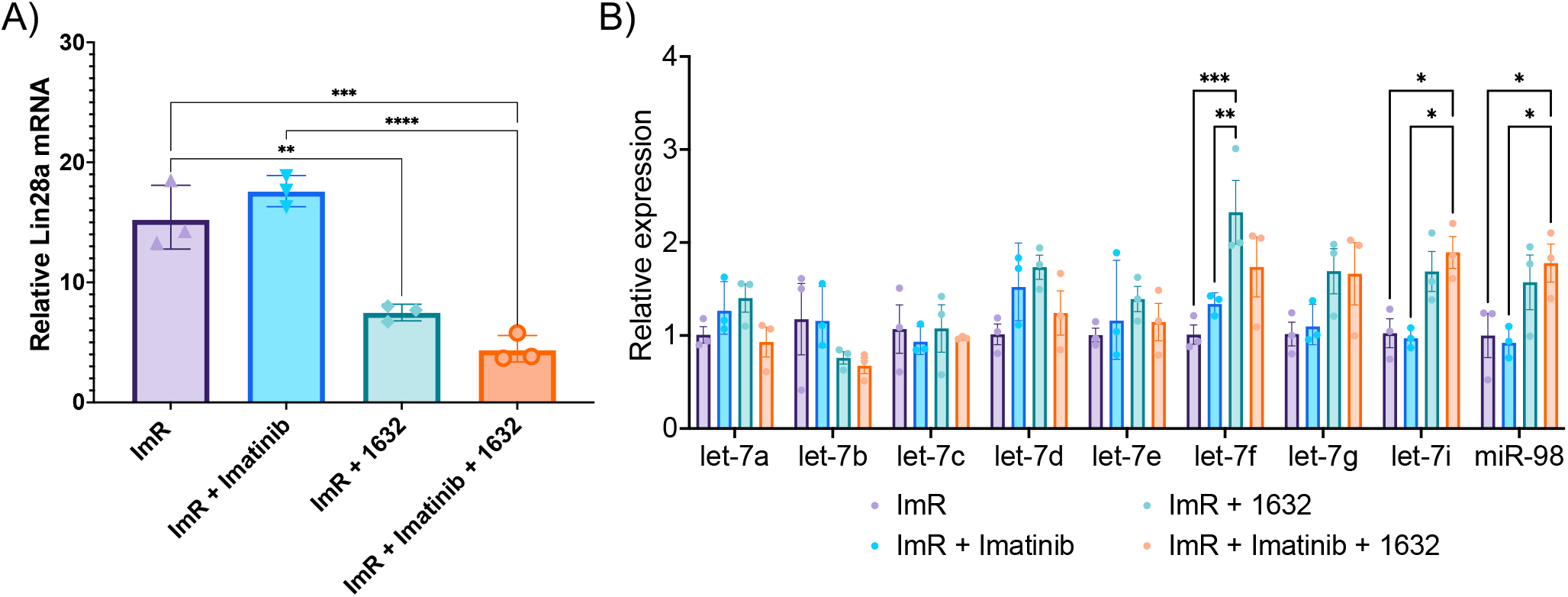
Synergistic combination of imatinib and 1632 alters the LIN28-Let-7 axis. (A) Relative abundance of Lin28a mRNA with the combination of imatinib and LIN28i-1632. (B) Comparison of let-7 family miRNA with the synergistic combination of imatinib and LIN28i-1632. Values were graphed as mean±SEM and statistical significance was determined by ANOVA test: *P <0.05; **P < 0.01; ***P < 0.001; ****P < 0.0001.

Inhibitors targeting the let-7-dependent axis (LI-71 and TPEN) showed no significant benefit. LI-71 (cold shock domain inhibitor) and TPEN (zinc-knuckle domain inhibitor) resulted in overall antagonistic or negligible synergy scores (-6.95 and -1.36, respectively) and failed to reduce cell counts compared to imatinib alone (Figure 1A-D). These results support a let-7-independent mechanism of resistance in this model. In contrast, LIN28i-1632 demonstrated strong synergism with imatinib (overall synergy score: 12.07), with peak synergy (29.71) at 50 μM LIN28i-1632 and 500 nM imatinib (Figure 1E). This combination reduced relative cell numbers by 71.15%, significantly outperforming either single treatment (Figure 1F). Based on these results, LIN28i-1632 was selected for further study.

### Temporal Proteomic Rewiring Over 48 Hours

To understand the dynamics of LIN28 inhibition, we performed time-course DIA proteomics over 48 hours of LIN28i-1632 treatment, quantifying 5,519 proteins. Peak proteomic remodelling occurred at 48 hours, with 458 differentially expressed proteins (DEPs) identified using FDR < 0.05 and >1.5-fold change (Figure S1A). An acute response to LIN28i-1632 occurred at 16 hours (36 DEPs) and 24 hours (133 DEPs). Most proteome-level changes occur between 0 and 48 hours (Figure S1B). Clustering analysis identified three primary response profiles: Cluster 1 (metabolic catabolism), Cluster 3 (RNA catalytic activity), and Cluster 4 (cytoskeletal regulation) (Figure S1D). Notably, the canonical LIN28 target HMGA1 and cell cycle regulators CDC123 and CDC20 were significantly downregulated at 48 hours (Figure S2). Conversely, GSEA identified an enrichment of G2M checkpoint proteins, mirroring the downregulation of pathways involved in oxidative phosphorylation and fatty acid metabolism previously linked to ImR resistance.

### Synergistic Treatment Promotes Apoptosis and G2/M Arrest

Using TMT-based proteomics, we analyzed the synergistic effect of the combination of imatinib and LIN28i-1632 at the 48-hour peak (Figure 3A). We observed 331 significantly changed proteins with the proteome using ANOVA with a minimum of 1.3-fold change and 0.05 FDR (Figure 3B). This combination treatment significantly upregulated proteins associated with interferon responses and apoptosis (CASP3, CASP6, BAX, PDCD4, and DIABLO), while downregulating mTOR signalling, MYC targets, and G2M checkpoint factors (Figures 3C and S3).

**Figure 3:**
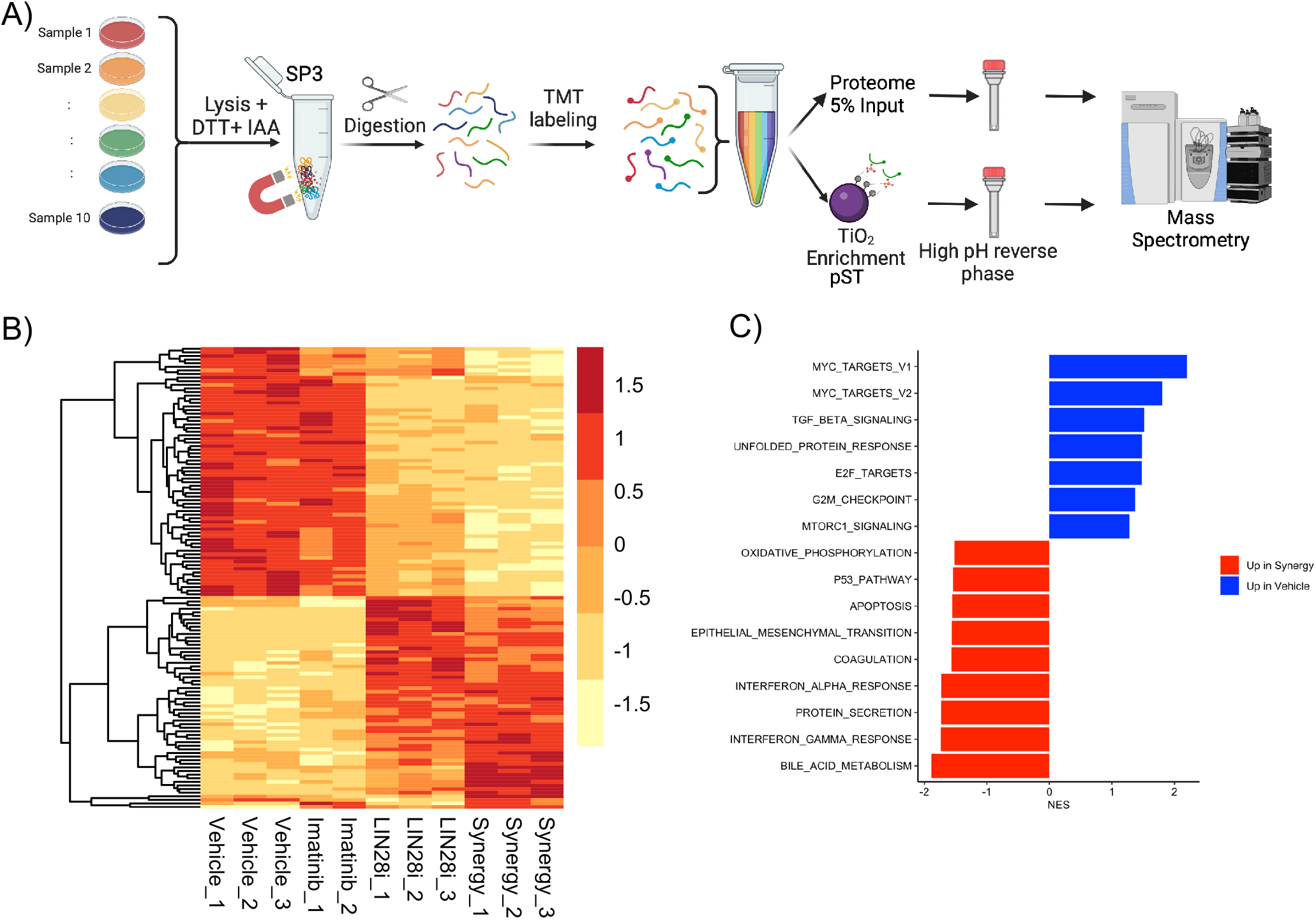
LIN28i-1632 primes the proteome for Imatinib sensitivity. (A) Mass spectrometry workflow showing the preparation of proteomics and phosphoproteomics samples. Heatmaps showing changes between proteome (A) and phosphoproteome (B) with greater than 0.3-log_2_ fold change and less than 0.05 FDR. (C) GSEA of Hallmark gene set comparing the vehicle and synergy of the proteome showing changes towards Myc targets, G_2_M and apoptosis.

We observed a significant decrease in the anti-apoptotic factor and G2 checkpoint protein, BCL2L1^33^ (Figure 4A). Functional validation confirmed these findings: Annexin V and Propidium Iodide (PI) staining revealed that the combination increased early apoptosis to 47.83%, compared to ∼10% for single treatments (Figure 4B). Additional, proteins involved in cell cycle regulation including MKI67, PBK, CUL4A, CDC45, TOP2A, and SRSF2 were downregulated (Figures 3C,4C and S4). Furthermore, flow cytometry showed that LIN28i-1632, both alone and with imatinib, induced an accumulation of cells in the G2 phase (34.0%) with a corresponding decrease in the S phase (Figure 4D).

**Figure 4:**
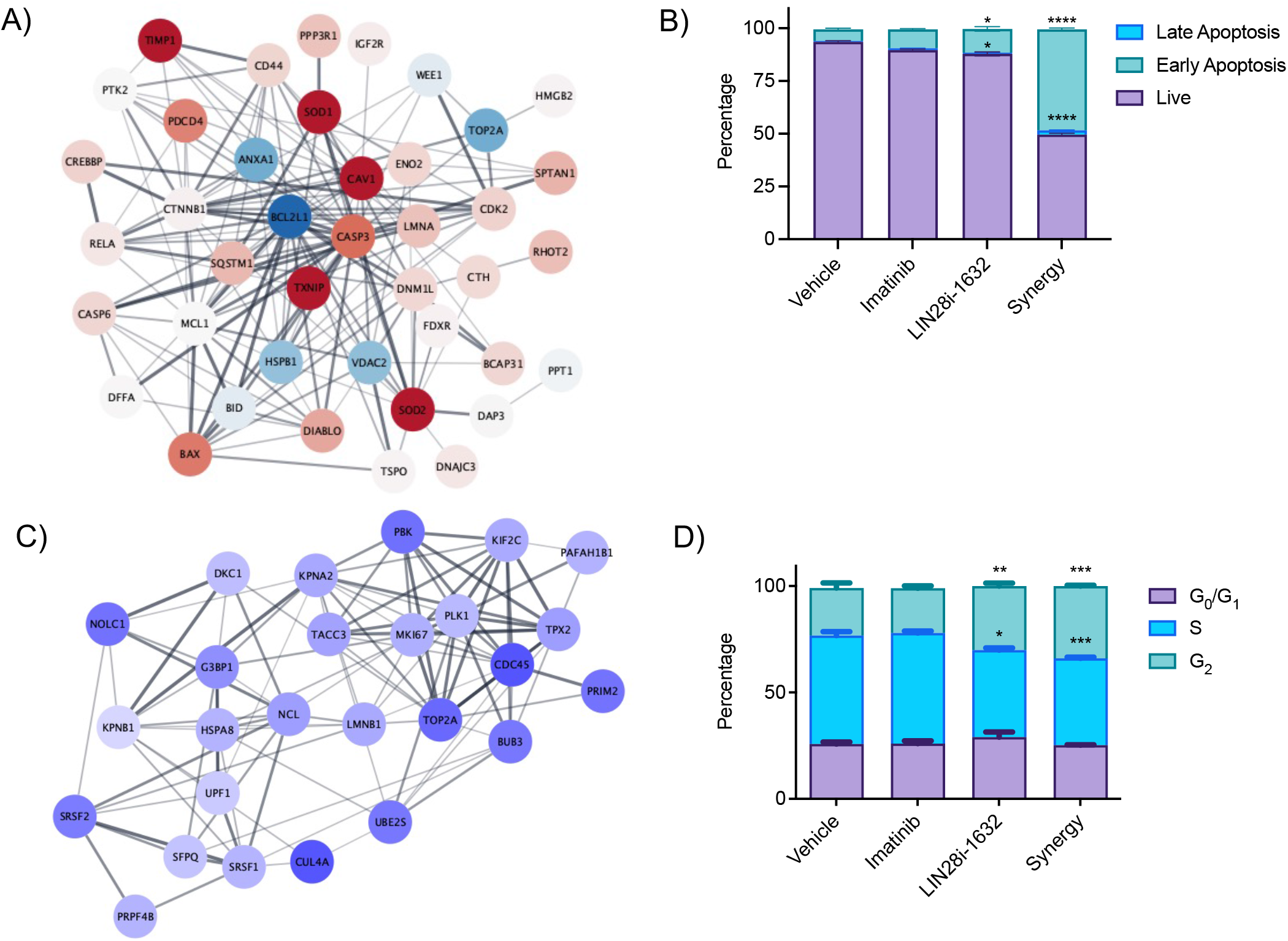
Synergy combination leads to changes in apoptosis and cell cycle. (A) Network analysis of genes associated with apoptosis hallmark gene set, comparing vehicle and synergy, genes higher in synergy in red and genes higher in vehicle in blue. (B) Annexin V and PI staining to assess percentage of live, early apoptosis and late apoptosis of ImR cells treated with either imatinib, LIN28i-1632 or synergistic combination (n=3, mean±SEM). (C) Network of gene found in G2M checkpoint hallmark gene set, showing genes higher in the vehicle in blue. (D) Analysis of cell cycle of ImR cells with either imatinib, LIN28i-1632 or synergistic combination by flow cytometry (n=3, mean±SEM). Statistical significance was determined by ANOVA test: *P <0.05; **P < 0.01; ***P < 0.001; ****P < 0.0001.

### Suppression of BCR-ABL and the AKT/RPS6K Axis

Using phosphoproteomics, we observed change in 55 phosphosites using ANOVA with a minimum of 1.3-fold change and 0.05 FDR (Figure 5A). Phosphoproteomic analysis using KSEA revealed that the synergistic combination significantly suppressed the activity of CDK1, AKT1, RPS6KA1, and RPS6KB1 (Figure 5B). Network analysis confirmed a decrease in the phosphorylation of key survival nodes, including GSK3A (pS21) and RPS6 (pS236) (Figure 5C).

**Figure 5:**
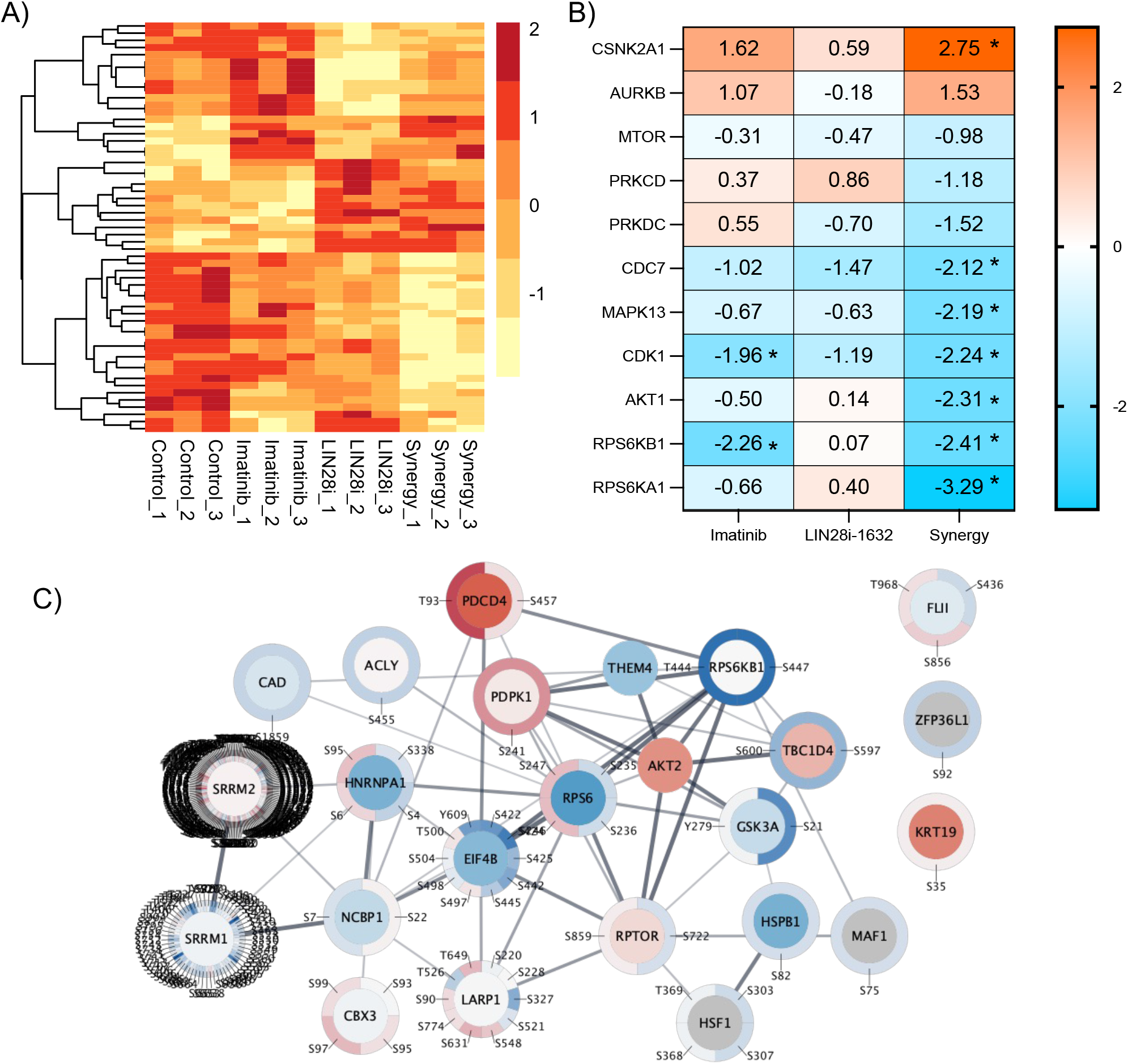
The combination of LIN28i-1632 and imatinib leads to changes in AKT and RPS6K signalling. (A) with greater than 0.3-log_2_ fold change and less than 0.05 FDR. (B) Kinase substrate enrichment analysis showing changes compared to vehicle control imatinib, LIN28i-1632 and synergy combination. KSEA scores with FDR <0.05 were noted with *. (C) Network of RPS6KA1, RPS6KB1, AKT and their substrates comparing vehicle and synergy combination with differential abundance (inner circle) and phosphorylation (outer circle).

Western blotting validated these proteomic trends. Unexpectedly, LIN28i-1632 treatment (alone or combined) led to a ∼2-fold decrease in BCR-ABL1 protein abundance without altering ABL1 levels (Figure 6A). Furthermore, the combination treatment restored PTEN levels (1.41-fold increase) and significantly reduced RICTOR abundance and AKT pS473 (3.21-fold decrease). Collectively, these results indicate that LIN28i-1632 restores imatinib sensitivity by simultaneously reducing BCR-ABL abundance and suppressing the AKT/RPS6K survival axis.

**Figure 6:**
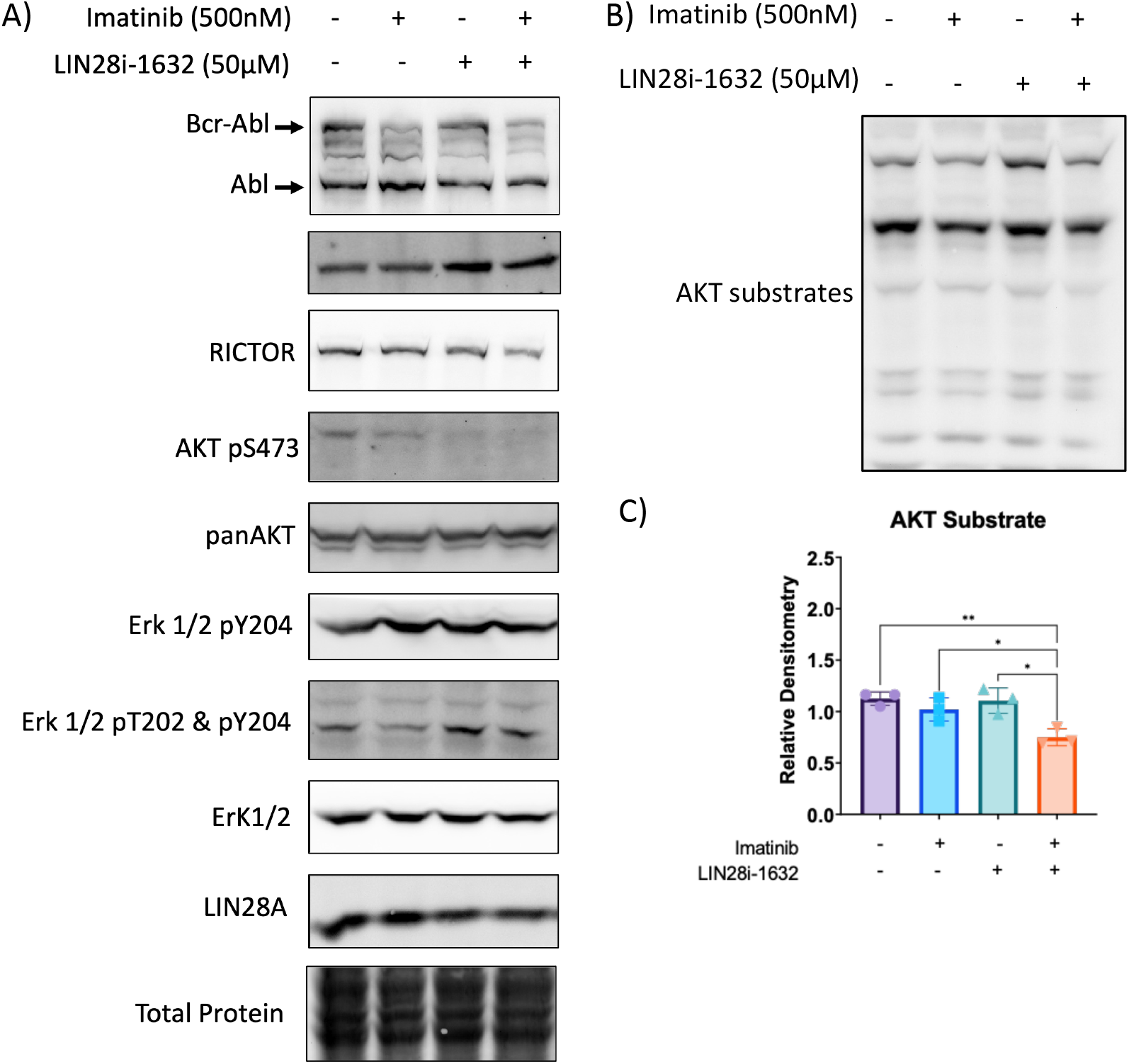
Combination of 1632 alters AKT signalling pathways. (A) Representative western blots of ABL, pY204 MAPK, pT202 & pY204 MAPK, MAPK, LIN28, PTEN, RICTOR, pS473 AKT, panAKT and total protein staining with imatinib, LIN28i-1632 and synergistic combination. Representative staining of AKT substrate blot (B) and corresponding densitometry (C). Statistical significance was determined by ANOVA test: *P <0.05; **P < 0.01.

## Discussion

The development of resistance to tyrosine kinase inhibitors (TKIs) like imatinib remains a primary hurdle in CML management, particularly when independent of BCR-ABL1 kinase domain mutations. Having previously identified LIN28A as a driver of this resistance, we explored whether pharmacological inhibition could restore sensitivity. Our screening of three inhibitors provided a critical mechanistic insight: while the let-7-dependent inhibitors (LI-71 and TPEN) failed to sensitize ImR cells, LIN28i-1632 showed potent synergy.

This suggests that imatinib resistance in our model is primarily driven by LIN28A’s let-7-independent functions ^16^. While LI-71 specifically disrupts the cold shock domain’s interaction with pre-let-7, LIN28i-1632 appears to possess a broader spectrum of activity, likely interfering with LIN28’s ability to directly bind and enhance the translation of oncogenic mRNAs. On the other hand, TPEN is a chelator with a high affinity for zinc and targets the zinc finger domain of LIN28; off-target effects on other zinc finger domains are highly likely ^13^. Unlike the previous two, the mechanism of inhibition of LIN28i-1632 has yet to be characterized ^14^. Intriguingly, LIN28i-1632, although less potent than LI-71, emerged as a promising candidate by effectively targeting both the LIN28-let-7 axis and let-7 independent functions, suggesting a broader spectrum of activity ^16^. Oral administration of LIN28i-1632 in mice was detected at the highest in blood, lung and liver, followed by the kidney, heart, and spleen ^34^. With negligible amounts detected in the brain tissue. Additionally, mice treated in two independent studies have had no effect on body weight while reducing tumour size. For these reasons, we choose to continue the investigation into LIN28i-1632.

Our DIA time-course analysis revealed that the cellular response to LIN28i-1632 is a progressive remodelling event towards 48 hours. Further timepoints are required to determine if further remodelling occurs at later timepoints. The early downregulation of HMGA1, a canonical LIN28 target, serves as the molecular “read-out” of successful LIN28 inhibition. However, the subsequent downregulation of CDC20 and CDC123 suggests a deeper impact on the proliferative capacity of CML cells. The observed G2/M arrest, indicates that LIN28 inhibition traps resistant cells in a vulnerable state, preventing the rapid cycling characteristic of TKI-resistant clones.

The most striking finding of this study is the dual-action mechanism of the LIN28i-1632/imatinib combination. While imatinib targets the activity of BCR-ABL, our data shows that LIN28i-1632 reduces the abundance of the BCR-ABL protein itself. This suggests that LIN28A may be involved in the translational maintenance of the fusion protein, a finding that warrants further investigation using CLIP-seq to determine if BCR-ABL1 mRNA is a direct LIN28 target or a different mechanism is involved.

Furthermore, our phosphoproteome and immunoblot data demonstrates that this combination effectively “short-circuits” the AKT/RPS6K survival axis. By increasing PTEN abundance and decreasing RICTOR levels, the combination treatment suppresses the PI3K/AKT pathway at multiple nodes. This is significant because AKT signalling is a well-documented escape route for CML cells under TKI pressure. Inhibition of LIN28 by LIN28i-1632 restored PTEN expression, likely by relieving the direct suppressive effect of LIN28 on PTEN translation. This restoration provides a secondary layer of signaling control, effectively blocking the PI3K/AKT survival bypass that typically allows cells to evade imatinib-mediated BCR-ABL inhibition.

While LIN28i-1632 shows promise, its off-target profile must be taken into account. Previous studies have noted potential binding to focal adhesion kinase (FAK/PTK2), which might contribute to the cytoskeletal and adhesion changes observed in the proteomic clusters. However, since FAK signalling also contributes to leukemia cell homing and niche-mediated resistance, this “off-target” effect may be therapeutically synergistic in a bone marrow environment ^15,35^

The temporal nature of the response suggests that the order of administration could be vital. Sequential therapy using a LIN28 inhibitor to “prime” cells by reducing BCR-ABL levels and slowing the cell cycle before administering a high-dose TKI could minimize toxicity and prevent the emergence of new mutations ^36,37^. Screening for analogs and employing computational modelling represent promising avenues for refining LIN28 inhibitors. Such efforts must improve specificity and binding kinetics while maintaining both let-7 and let-7 independent functions of LIN28i-1632 to treat cancer resistance.

This study establishes LIN28i-1632 as a potential tool for overcoming TKI resistance in CML. By simultaneously depleting the BCR-ABL protein reservoir and silencing the AKT/mTOR survival axis, the combination treatment drives resistant cells towards apoptosis.

These findings support the development of next-generation LIN28 inhibitors as a viable strategy for patients who have exhausted standard TKI options.

## ACKNOWLEDGMENTS

We thank Kun Ping Lu, Xiao Zhen Zhou, and Patrick O’Donoghue for their valuable insights and critical discussions. This research was supported by grants from the Canadian Institute of Health Research (to SSCL), the Natural Sciences and Engineering Research Council of Canada (to SSCL and IUH); the Ontario Ministry of Research and Innovation (to IUH). OFJH received funding from Thermo Scientific Tandem Mass Tag Research Award, which provided reagents for this project. AW and MIF each held a Queen Elizabeth II Graduate Scholarship in Science and Technology (QEII-GSST). SSCL holds the Canada Research Chair and Wolfe Medical Research Professorship in Molecular and Epigenetic Basis of Cancer.

## AUTHOR CONTRIBUTIONS

SSCL and OFJH conceived and designed the project. OFJH, TW, AW and JHK performed the research. OFJH and TW analyzed the data. SSCL and OFJH wrote the manuscript with input from AW, MIF and IUH.

