## Supplementary Figures for "Pharmacological inhibition of LIN28A promotes imatinib sensitivity in CML resistance"

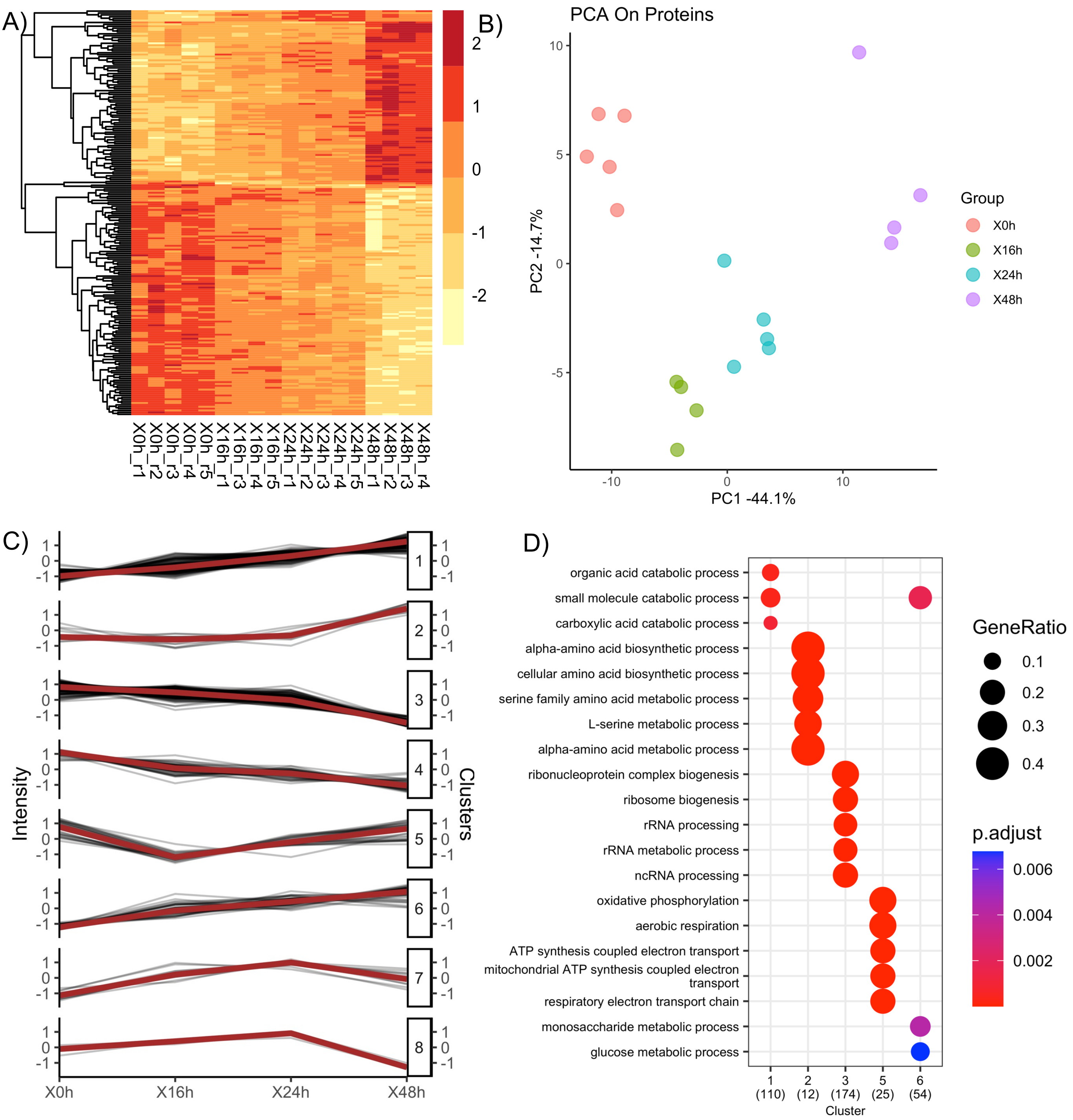


**Figure S1: Time course of LIN28i-1632 treatment showing translational rewiring**. (A) Unsupervised clustering of PCA showing separation of time across the first component. (B) Heatmap showing significantly differently regulated (C) Showing the eight clusters profile plots optimized from gap statistic. (D) Gene ontology enrichment of biological processes derived from the clusters of significant features.

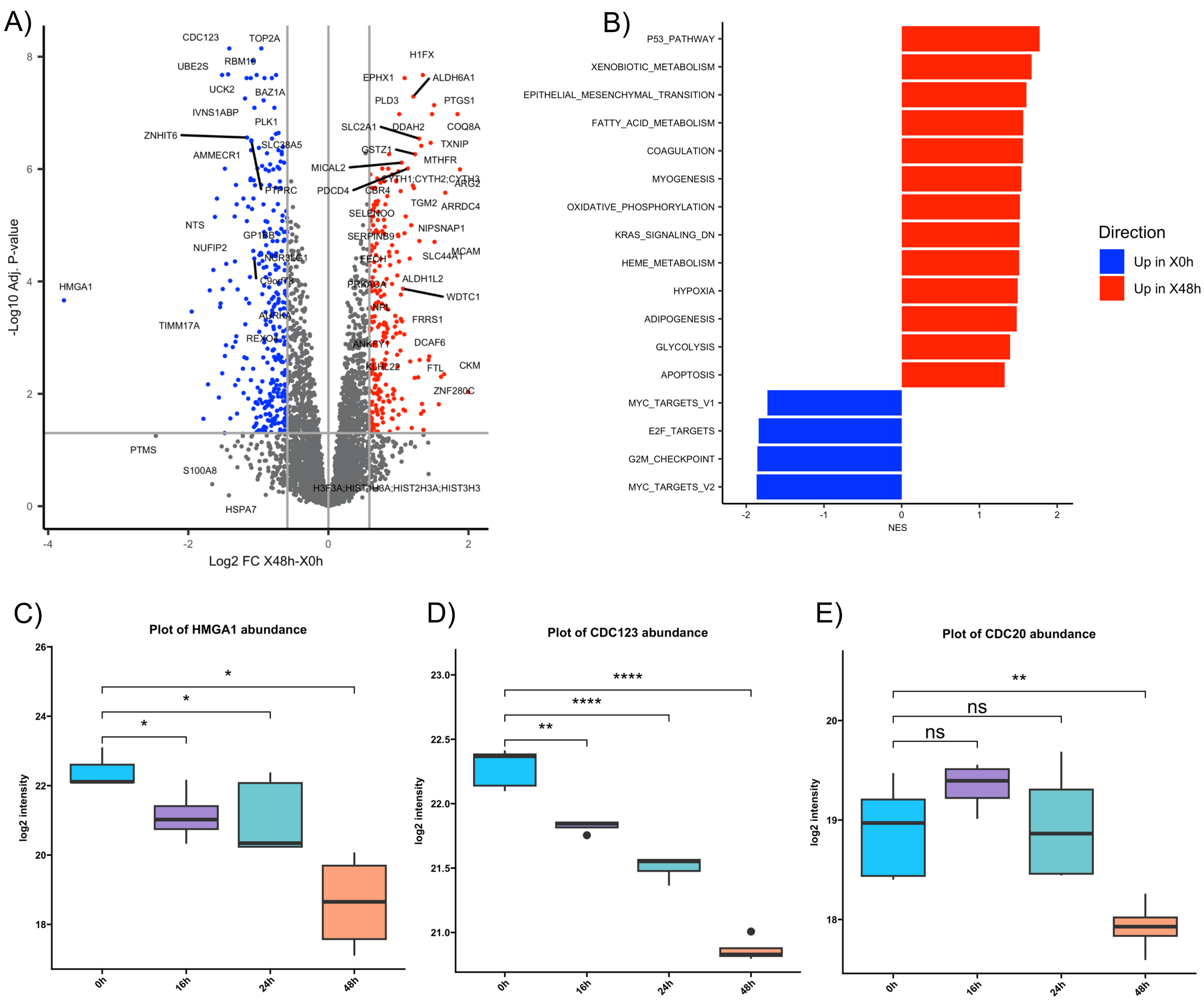


**Figure S2: Metabolism, cell cycle and stem cell associated genes are alternated at 48 hours of LIN28i-1632 treatment**. (A) Volcano plot showing significant changes comparing 0h timepoint and 48h time points treated with LIN28i-1632 with genes highlighted with FDR less than 0.05 and log2-fold change greater than 0.5 fold change. (B) GSEA of Hallmark gene set comparing the 0 and 48hs of the proteome. Relative abundance comparing vehicle, imatinib, LIN28i-1632 and synergy combination of HMGA1 (A), CDC123 (B), CDC20.

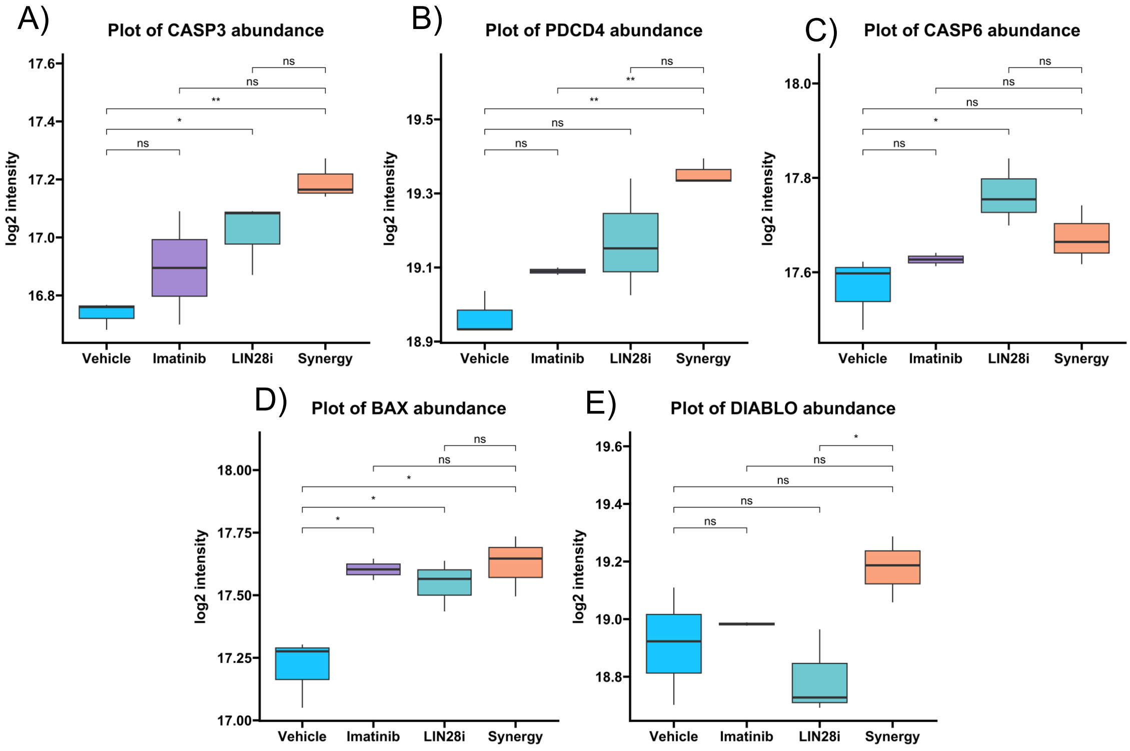


**Figure S3: Relative protein abundance of select proteins involved in apoptosis in reference to figure 4.** Relative abundance comparing vehicle, imatinib, LIN28i-1632 and synergy combination of CASP3 (A), PDCD4 (B), CASP6 (C), BAX (D) and DIABLO (E).

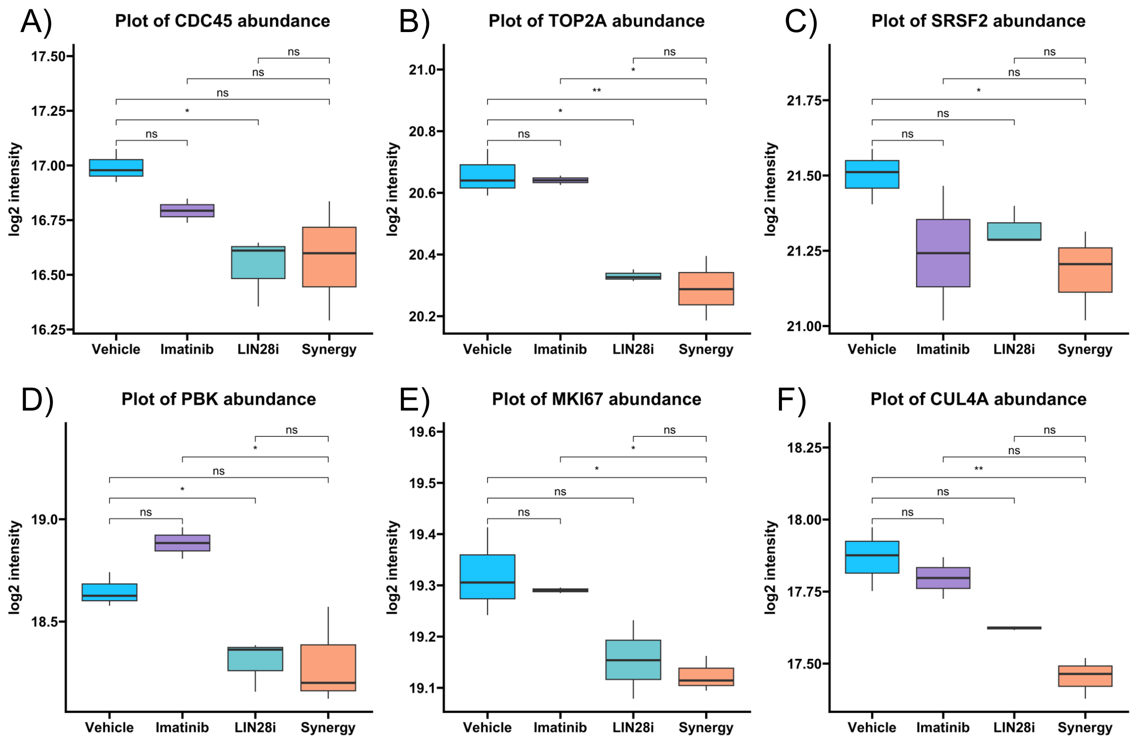


**Figure S4: Relative protein abundance of select proteins involved in G2M checkpoint in reference to figure 4.** Relative abundance comparing vehicle, imatinib, LIN28i-1632 and synergy combination of CDC45 (A), TOP2A (B), SRSF2 (C), PBK (D), MKI67 (E) and CUL4A (F).

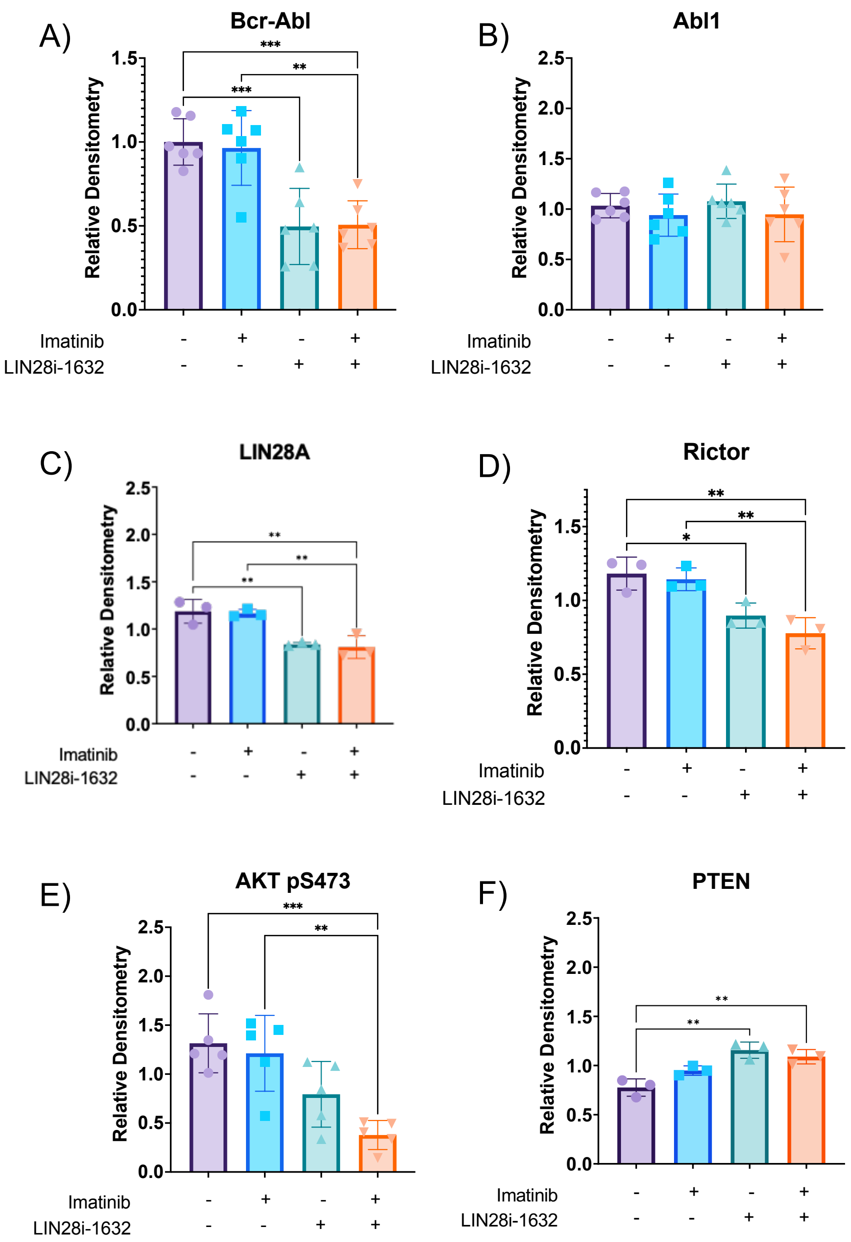


**Figure S5: Densitometry of western blots highlighting changes in AKT signalling in relation to Figure 6**. (A-F) Comparison of vehicle control, single treatment of imatinib and LIN28i-1632, and the synergistic combination densitometry from western blots in figure 5 showing BCR-ABL (A), ABL1 (B), LIN28A (C), RICTOR (D), AKT pS473 (E) and PTEN (F).
